# Capturing the biology of disease severity in an organoid model of LIS1-lissencephaly

**DOI:** 10.1101/2022.12.19.520907

**Authors:** Lea Zillich, Matteo Gasparotto, Andrea Carlo Rossetti, Olivia Fechtner, Camille Maillard, Anne Hoffrichter, Eric Zillich, Ammar Jabali, Fabio Marsoner, Annasara Artioli, Ruven Wilkens, Christina B. Schroeter, Andreas Hentschel, Stephanie H. Witt, Nico Melzer, Sven G. Meuth, Tobias Ruck, Philipp Koch, Andreas Roos, Nadia Bahi-Buisson, Fiona Francis, Julia Ladewig

## Abstract

Lissencephaly is a malformation of cortical development (MCD) characterized by reduced to absent gyri and a disorganized cortex, leading to severe neurological consequences in affected individuals, including epilepsy, intellectual disability, and reduced life expectancy. Treatments are purely symptomatic, and patients often remain refractory to them. Heterozygous mutations in the *LIS1* gene, encoding a regulator of the microtubule motor dynein, cause LIS1-lissencephaly. For unknown reasons, LIS1-lissencephaly patients show marked differences in disease severity despite each carrying a heterozygous LIS1 mutation. We leveraged forebrain-type organoids from patients diagnosed with mild, moderate, or severe LIS1-lissencephaly to investigate, in a cytoarchitecture and multi-omics approach, disease and severity grade associated phenotypes, mechanisms, and rescue approaches. We identified alterations of the cytoarchitecture, progenitor cell homeostasis, and neurogenesis often with a severity-dependent gradient. Identified disease-linked molecular mechanisms were microtubule destabilization, WNT-signaling, protein metabolism, and perturbed cadherin- and unfolded protein-binding. Some mechanisms exhibited a severity-dependent gradient or were specific to a severe grade. We present strategies to reverse phenotypic changes in LIS1-patient organoids and identify mTOR pathway inhibitors in *in silico* drug repurposing analysis as potential novel therapeutic strategy. By probing the top hit drug, the mTOR inhibitor everolimus, we could indeed rescue severity-dependent phenotypic changes in LIS1-patient organoids. This study demonstrates that organoid-based modeling is sensitive in recapitulating disease severity, which presents an important step in patient stratification, and allows the development of novel personalized rescue strategies with therapeutic potential.

## Introduction

The human neocortex, critical for language, sociability, and sensorimotor control, is an expanded, highly organized, and extensively folded, gyrencephalic structure (1). Malformations of human cortical development (MCD) are a vast and heterogeneous group of disorders with genetic and/or environmental etiology, often associated with disruption of cerebral cortex architecture. They account for up to 40% of childhood treatment-refractory epilepsies (2).

Heterozygous mutations in the *LIS1* gene cause lissencephaly (smooth brain; MIM: 607432) in humans with diverse neuroimaging phenotypes, ranging from mild pachygyria (broad gyri) to severe agyria (no gyri), resulting in clinical phenotypes including treatment-refractory epilepsy and intellectual disabilities (3). While the clinical disease spectrum correlates with the degree of lissencephaly, location and type of mutation do not (4). The *LIS1* gene encodes a protein that plays a functional role in dynein-dynactin assembly and orchestrates microtubule motor function vital for cellular organization and division (5, 6). Studies on Lis1-mouse models unveiled that LIS1-lissencephaly is likely to be associated with perturbance in dynein-dependent processes such as neuronal migration and progenitor cell behavior including interkinetic nuclear migration and mitotic spindle orientation (7-11). Although the phenotypes observed were dramatically milder in mouse models - which are inherently lissencephalic - than in humans, these studies suggest that Lis1 protein dosage is relevant to phenotypic severity (7-10).

Due to limited access to human postmortem brain tissue from LIS1-lissencephaly patients (LIS1-patients) at different developmental stages, it is difficult to study disease, and severity-related molecular and cellular drivers in the human context. *In vitro* disease modeling using induced pluripotent stem (iPS) cell-derived cerebral organoids combined with advances in OMICS technologies have shown considerable potential to understand human-specific processes in brain development and disease (12). Organoids are particularly powerful, especially for organs that have undergone large evolutionary changes, such as the human brain and for disorders with developmental origin where access to affected primary tissue is limited. Furthermore, the technology allows a personalized approach to disease biology and treatment. Exploiting organoids, we and others modeled a very severe type of lissencephaly, the Miller-Dieker-Syndrome (MDS, MIM: 247200), a contiguous gene deletion syndrome including *LIS1* (13-15). These studies suggest a role of LIS1 in neurogenesis and progenitor cell mitosis and propose how LIS1 deficiency may result in an impairment of ventricular niche signaling and cell fate control via the N-cadherin/ß-catenin signaling axis (13). More recently, organoids also allowed us to describe a role of LIS1 in a newly discovered dynein motor-dependent movement of human basal radial glia (bRG) cells during interphase, called interphasic somal translocation (IST), contributing to bRG cell dissemination in the human cortex (16).

While there has been considerable success in identifying certain developmental aspects related to lissencephaly using iPS cell-derived organoids, the ability to recapitulate disease severity in a defined patient cohort has not yet been demonstrated. Here, we use LIS1-lissencephaly as a platform to address whether organoid technology is sufficiently sensitive to predict disease severity among individual patients, and can enhance our understanding of disease mechanisms and by that, contribute to the development of novel therapeutic strategies.

## Results

### Modeling mild, moderate, and severe LIS1-lissencephaly in patient-derived cerebral organoids identifies severity-dependent changes in premature neurogenesis and progenitor cell homeostasis

To model disease severity of LIS1-lissencephaly, we selected seven LIS1-patients covering a spectrum of gyrification alterations (Fig. 1A), ranging from Dobyns grade 5 (mild) to 1 (severe) from a cohort comprising 63 cases (17, 18) (for more details on MRI data see Philbert, Maillard (19); for selection criteria see Material and Methods section). Each patient harbors a molecularly characterized heterozygous pathogenic variant in the *LIS1* gene (Sup. Fig. S1A). Following the reprogramming of patient-derived somatic cells to iPS cells, and their basic characterization (2 clones each, Sup. Fig. S1B-G, Sup. Table S1, we validated the respective patient-specific *LIS1* mutations by sequencing (Sup. Fig. S1H-J). We selected seven iPS cell lines from a repertoire of healthy controls, including six from early age comparable to the LIS1-patients, and one from middle age (Sup. Table S1).

**Fig. 1.**
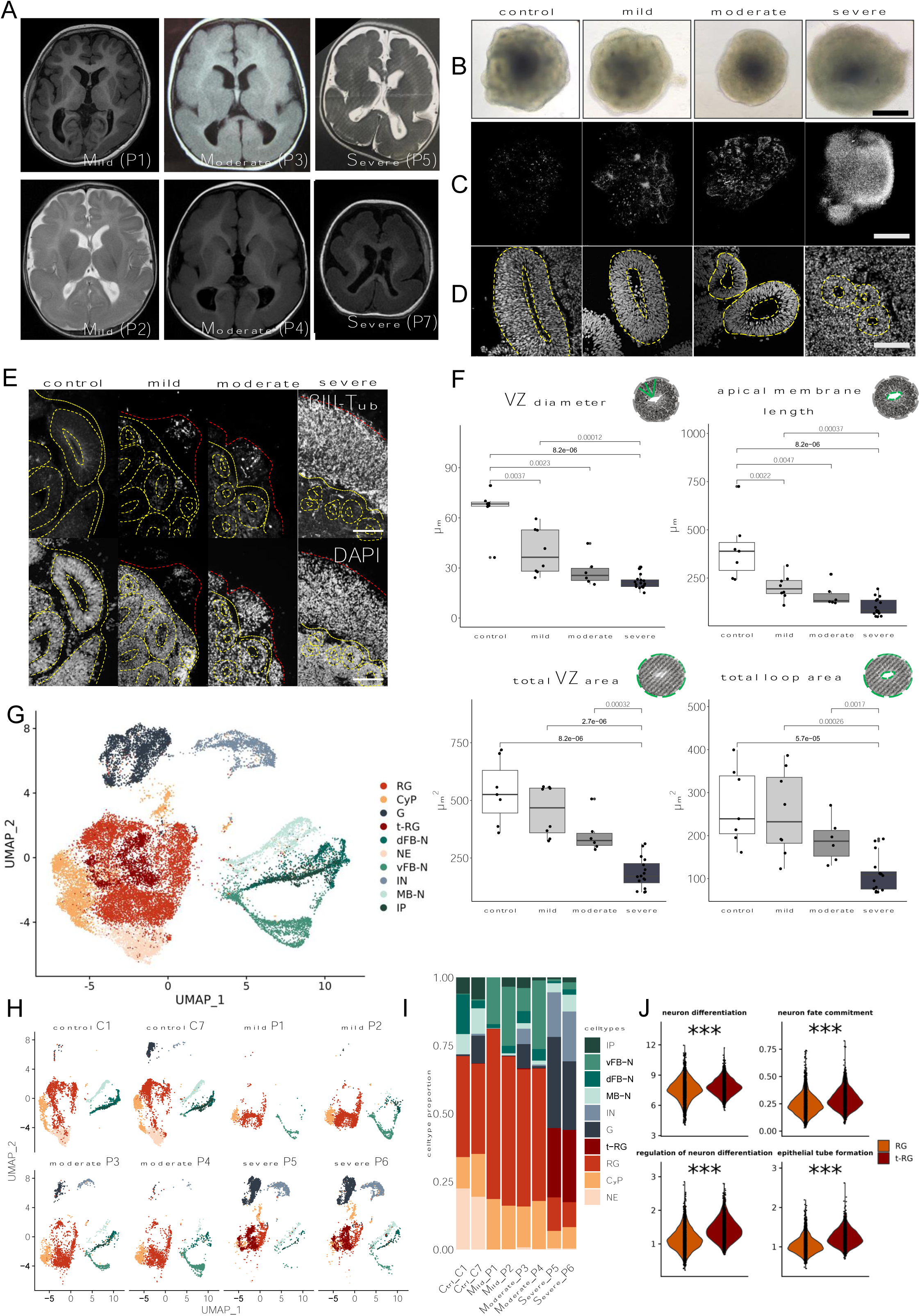
Morphological and cellular characterization of LIS1 patient-derived organoids across severity grades. **A**) LIS1-patient MRIs for mild, moderate and severe LIS1-lissencephaly. LIS1-mild grade patients 1 and 2 exhibit a posterior pachygyria (smooth cortex) with subnormal frontal gyration. LIS1-moderate grade patients 3 and 4 show a posterior pachygyria in the parietal and occipital lobes, and mild frontal pachygyria. LIS1-severe grade patient 5 and 7 exhibit a diffuse pachygyria, almost complete agyria predominantly in the parietal and occipital lobes (with a sparse cell layer). Personal information from the original MRI of patient 5 has been blackened in the upper left corner. **B**) Representative brightfield images of control, mild, moderate and severe LIS1 patient-derived organoids at day 20. **C**) Representative light sheet microscopy (LSM) pictures of whole-tissue cleared control, mild, moderate and severe patient-derived organoids at day 20, stained for ß-III Tubulin (TUBB3). **D**) Representative Hoechst staining of ventricular zone structures (VZ) of control, mild, moderate and severe patient-derived organoids at day 20. Yellow dotted lines define the VZ areas. **E**) Representative beta-III-tubulin (TUBB3) and DAPI counterstained pictures of control, mild, moderate and severe LIS1-patient derived organoids at day 20. **F**) Quantification of structural VZ parameters in control and LIS1 patient-derived organoids at day 20. Central line in boxplot represents median, lower lines the 25^th^ and upper the 75^th^ percentile, Whiskers are 1.5 interquartile ranges. Individual dots represent the mean of one batch. **G)** UMAP dimensional reduction and unbiased clustering reveals 10 distinct color-coded cell populations within control, mild-, moderate- and severe LIS1 patient-derived organoids: neuroepithelial cells (NE), cycling progenitors (CyP), radial glia cells (RG), intermediate progenitors (IP), transitory RG (t-RG), dorsal forebrain (dFB-N), ventral forebrain (vFB-N), midbrain (MB-N), interneurons (IN) and glial cells (G) **H**) UMAP graphs split by samples. Number of organoids and cells analyzed: 23 organoids, control N=6, mild N=6, moderate N=5, severe N=6 (two independent cell lines per condition: 2 controls (8647 single cells), 2 mild patients (3444 single cells), 2 moderate patients (5168 single cells) and 2 severe patients (7723 single cells). **I)** Cell type distribution in each severity grade divided by sample. **J)** Violin plots depicting scores of GO terms differentiating t-RG from RG cells, ***P< 0.001. Scalebars: B, C 200 µm; D, E 50 µm.

We then generated forebrain-type cerebral organoids (20) from the seven LIS1-patients and seven control iPS cell lines. While organoids from controls and patients with a milder malformation gradually developed regular neuroepithelial loop-like structures, organoids from patients with a moderate disease grade appeared generally smaller in size (Fig. 1B). Organoids from severely affected patients were not smaller compared to control-derived organoids but developed irregular edges, with single cells noticeably growing out from the structures (Sup. Fig. S2A). Immunohistochemical analyses following whole-tissue clearing or cryo-sectioning showed that all successfully generated organoids from severe malformation patients presented a large belt of neurons which was less abundant in mild and moderate conditions and nearly absent in control-derived organoids at this age (Fig. 1C-E). To address whether the LIS1 organoid model can capture differences between control, mild, moderate, and severe LIS1-lissencephaly, we assessed the architecture of the individual VZ-like structures within the organoids by analyzing multiple VZ morphological parameters, i.e. the apical and basal membrane lengths, the VZ diameter and size as well as the total loop area and size (13). Of the six parameters analyzed, we found significant differences in all parameters in organoids derived from patients with severe, four parameters in moderate, and two in mild severity grades, compared to control organoids. Between patient samples, we found that mild and severe separated in five out of six parameters and moderate and severe in three out of the six parameters analyzed (Fig. 1F, Sup. Fig. S2B, Sup. Table S2). Overall, our data point towards a severity-dependent gradient in the VZ morphological parameters from mild, moderate to severe LIS1-patient organoids.

To further investigate changes in cell type composition between control, mild, moderate and severe LIS1-patient organoids, and the underlying molecular mechanisms of the severity-dependent gradient in phenotypic changes, we profiled organoids by single-cell RNA sequencing (scRNA-seq; two to three pooled organoids at day 23±2 from two different donors per condition; Fig. 1G). We observed high homogeneity in cell type composition within each severity grade, independent of the patient-specific genetic backgrounds (Fig. 1H-I). We identified nine cell populations based on known marker genes including neuroepithelial cells (NE), cycling progenitors (CyP), radial glia cells (RG), intermediate progenitors (IP), dorsal forebrain (dFB-N), ventral forebrain (vFB-N), midbrain (MB-N) and inter-neurons (IN), as well as astroglia cells (G) (21-25) (Sup. Fig. S3A). We further identified a progenitor cell cluster that exhibits a very similar marker profile as the RG population but clusters separately. GO term analyses revealed that this cell cluster distinguishes from the RG cluster by a significant overrepresentation of GO terms linked to neuronal differentiation, including generation of neurons, neuron fate commitment, and regulation of neuron differentiation, thus termed transitory RG (t-RG; Fig. 1J, Sup. Table S3). Pseudotime analysis confirmed the developmental trajectories of the cells, starting with NE cells and developing via RG types to neuronal cell types (Sup. Fig. S3B). A comparison of cell type proportions between the severity grades revealed a significant reduction in neural progenitor cells accompanied by a significant increase of differentiated cells in the LIS1 severe grade condition (Sup. Fig. S3C). We further investigated the progenitor cell type composition in more detail. We identified a depletion of neuroepithelial (NE) cells in all patient samples. While the mild and moderate grade LIS1-patient organoids showed a relative increase in RG cells, the samples derived from severe grade patients showed a clear reduction of this population compared to the control, mild and moderate condition, accompanied by the appearance of the severe grade-specific t-RG cluster. Taken together, the increased abundance of neuronal cells, the depletion of early-stage NE cells, the upregulation of the later-stage RG cells in mild and moderate, and of t-RG in the severe condition suggest that premature neurogenesis is likely to drive the change in cell type composition. To exclude that apoptosis does not lead to changes in progenitor cell type composition, we investigated the cumulative expression of genes in apoptosis-related GO terms in the scRNA-seq data and found no significant changes across the different genetic backgrounds and severity grades (Sup. Fig. S3D). Taken together, our morphological characterization and transcriptional analyses suggest common and severity-dependent alterations in progenitor cell homeostasis and neurogenesis.

### Multi-omics analyses suggest shared and severity-specific molecular alterations underlying the phenotypic changes across the different LIS1-severity grades

To elucidate common disease mechanisms and associated severity-dependent changes, we performed differential expression analysis in the pooled progenitor cell types (NE, RG, t-RG), excluding cycling progenitors to avoid confounding by cell cycle. This revealed 140 significantly differentially expressed genes in the mild, 80 in the moderate, and 104 in the severe condition compared to the control with specific and shared genes for each severity grade (Fig. 2A, Sup. Table S4-S6). We observed 57 overlapping DE genes between the mild and moderate conditions, 41 between moderate and severe, and 38 between mild and severe (Fig. 2B). The most significant, specific association with the mild condition is the upregulation of *FZD5*, coding for a receptor for Wnt5A involved in forebrain and cortical hem development (26). The long non-coding RNA 51, alreadyfound to be expressed in the human neocortex (27), and the pro-neurogenic gene *MEIS2*, a gene already associated with MCD (25), were found to be the most significant uniquely associated transcripts within the moderate condition. In the severe condition, the top associations were a downregulation in *MTRNR2L12*, a gene associated with MCD and synaptic connectivity (28, 29), and an upregulation in *NNAT*, a gene involved in brain development and in neuronal functionality (30, 31). Several genes were differentially expressed across conditions, such as *DLK1*, a transmembrane protein regulating cell growth, and involved in neuronal differentiation and homeostasis (32), *SFRP1*, a WNT pathway modulator (33), and *LHX2* as well as *FOXG1*, both of which play fundamental roles in early cortical specification (34, 35).

**Fig. 2.**
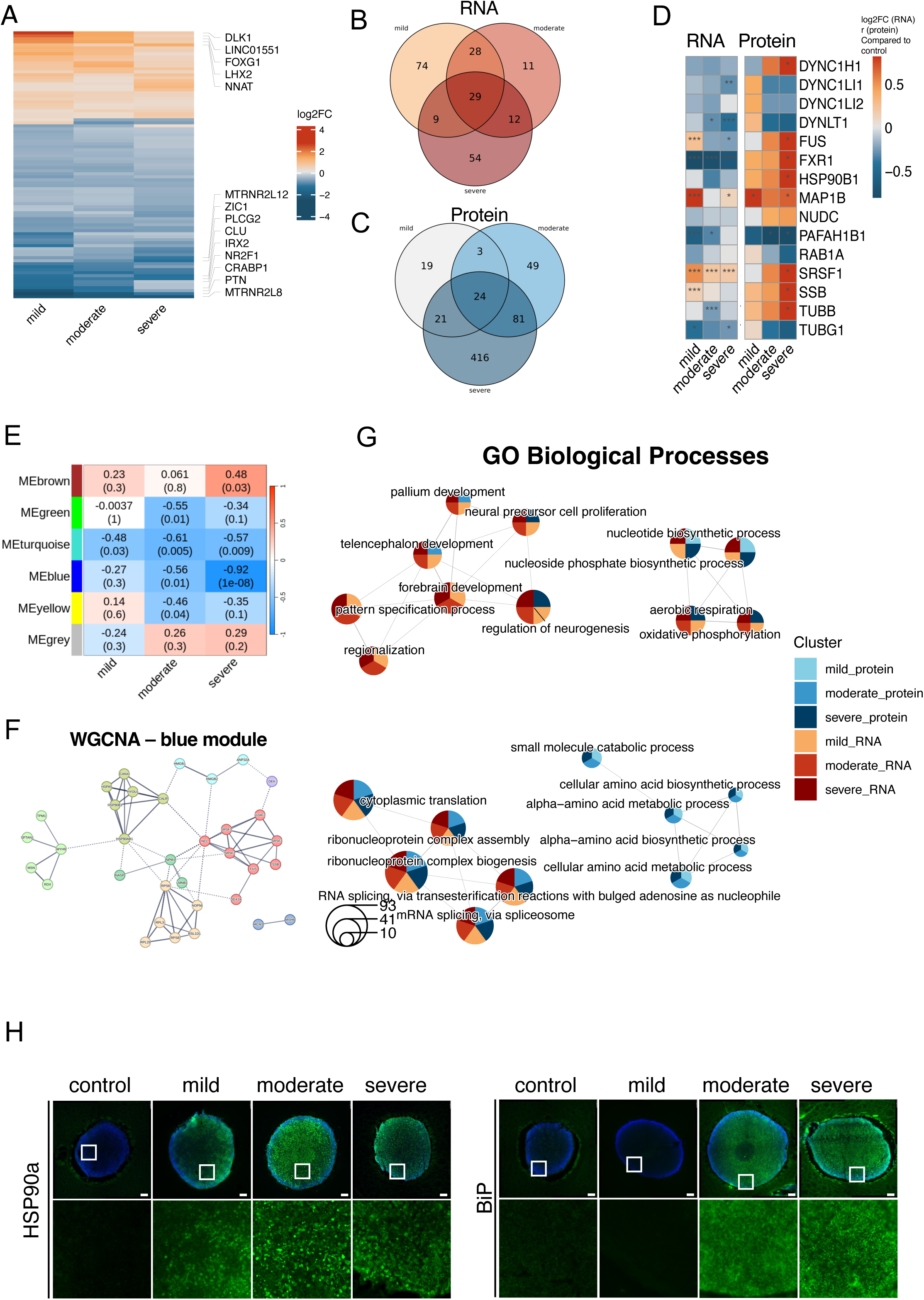
Transcriptomic and proteomic dysregulation in LIS1 patient-derived organoids across severity grades. **A)** Heatmap depicting differentially expressed genes between the control and the mild, moderate, and severe severity-grades (adjusted p-value < 0.05 and absolute log2FC > 0.50). Annotated are the 10 unique genes with the smallest p value per condition. Color intensity refers to the strength of association (log2FC). **B)** Venn diagram depicting the overlap between differentially expressed genes between the mild, moderate, severe and control conditions. **C)** Venn diagram depicting the overlap between differentially expressed proteins between the mild, moderate, severe and control conditions. **D)** Heatmap depicting log2FC (RNA) and Wilcoxon’s r (protein) between the severity grades and the control condition, for LIS1 interaction partners. **E)** Heatmap depicting the correlations of module eigengenes from WGCNA modules and severity grades. **F)** String network of proteins included in the blue module. Proteins have been clustered by MCL clustering with the inflation parameter set to 3. Edges represent the degree of confidence for the interaction and only proteins forming interactions with a high degree of confidence (interaction score ≥ 0.700) have been plotted. Red cluster: associated with mRNA splicing. Yellow cluster: involvement in cytoplasmic translation and rRNA processing. Sage green: involved in protein folding in the endoplasmic reticulum. Mint and dark green clusters: associated with the regulation of cell shape and chromatin assembly. **G)** Emap plot of GO biological processes overrepresented in differentially regulated genes and proteins, showing five categories, size of circles representing the number of differentially regulated features in the GO term. **H)** Representative widefield images of D18 forebrain organoids stained with HSP90a and BiP. Scalebar of overviews 100 µm; details 20 µm.

To further investigate the common and severity-dependent pathomechanisms of LIS1-lissencephaly at the protein level, we generated homogeneous cortical progenitor cells in 2D cultures according to established protocols from control, mild, moderate, and severe LIS1-patient-derived iPS cells (13, 36). Following basic characterization (Sup. Fig. S3F), we quantified protein abundances using label-free mass spectrometry. When comparing the proteomic signatures of the severity grades to the control condition, we observed 67 differentially regulated proteins in the mild, 157 in the moderate, and 542 in the severe grade patient-derived samples, although these findings were significant only before adjusting for multiple testing (Fig. 2C, Sup. Fig. S3E, Sup. Table S7).

As Lis1-mouse models suggest that Lis1-protein dosage correlates with the degree of phenotypic changes, we investigated LIS1 protein in the different conditions (control, mild, moderate, and severe). LIS1 (PAFAH1B1) protein levels were strongly decreased in a severity-dependent manner in progenitor cells. However, at the transcript level, LIS1 showed the strongest downregulation in mild severity cases, while no downregulation was observed in severe cases (Fig. 2D). We further investigated RNA expression and protein levels of known LIS1 interaction partners (e.g. FUS, SRSF1, GSP90, SSB, DYNC1H1, and TUBB) and identified that the progressive de-regulation of their protein expression correlates with the severity grade, only partially corresponding to transcript levels (Fig. 2D). We further performed a network analysis (WGCNA) of the protein data and identified five modules, two of them significantly associated with the LIS1-lissencephaly severity grades (Fig. 2E). The turquoise module showed significant downregulation in the mild, moderate and severe samples and GO enrichment analysis showed the inclusion of proteins involved in proteasomal protein degradation, ubiquitin-dependent protein catabolism, protein neddylation, cytoplasmic translation, and L-serine biosynthetic process (Sup. Table S8, Sup. Fig. S3G). Notably, several proteins involved in protein catabolism, including members of the proteasome and the COP9 signalosome, are found in this module, indicating that upregulation of protein catabolism is a feature shared by all LIS1-patient-derived organoids, regardless of the severity grade. In addition, the blue module showed an increasing severity grade-dependent negative correlation (Fig. 2E). GO enrichment analyses revealed that proteins belonging to this module were enriched amongst others for pathways related to RNA splicing, cadherin binding, G protein activity, and unfolded protein binding (UPB, Sup. Table S8). Network analysis of this module identified three major clusters (Fig. 2F).Notably, the first subset of proteins includes such involved in RNA splicing that have been previously identified as nuclear interactors of LIS1 in *AGO2* gene knockout embryonic stem cells (37). Among them, NCL and DXCX3 have also been associated with pre-rRNA transcription, ribosome assembly, and translation of components of the core translational machinery (38, 39). The second and third groups contain proteins involved in ribosome assembly and enhancers of protein translation and proteins involved in endoplasmic reticulum (ER) protein folding, including the ER chaperone BiP, a primary sensor in the activation of ER stress via upregulation of UPR (40) indicating that the misregulation of translational activity, already suggested by components of the turquoise module, may worsen in a severity-dependent fashion. The expression of BiP and the protein-folding chaperone HSP90a was confirmed by immunofluorescence in day 20 organoids. While HSP90 was upregulated in all severity grades, expression of BiP was enhanced in moderate and severe conditions (Fig. 2H).

### Integrative analysis of differential RNA and protein regulation identified critical severity grade-dependent phenotypes in LIS1-patient organoids

An integrative GO enrichment analysis was performed based on the results of the differential RNA and protein expression analyses for biological processes (Fig. 2G, full results Sup. Table S9) and molecular functions (Sup. Fig. S3H, full results Sup. Table S10). We found changes in GO clusters connected to early brain development including forebrain development, neural precursor cell proliferation, and regulation of neurogenesis, indicating early neurodevelopmental abnormalities in line with our morphological characterization of the patient-derived organoids. This cluster showed strong enrichment for differentially expressed genes across all severity grades, often with a severity-dependent gradient at the RNA level, and in moderate and severe conditions at the protein level. We also observed clusters with strong enrichment across all severity grades and -omics layers; one related to triphosphate metabolic processes and one related to RNA splicing and cytoplasmic translation, indicating broad changes in LIS1-lissencephaly independent of the severity grade (Sup. Table S9-S10). For molecular functions, we showed, besides others, enrichment for pathways related to tubulin-, actin-, cadherin-, and WNT-protein binding (Sup. Fig. S3H, Sup. Table S10).

When investigating GO term enrichment in more detail we identified amongst others a gradual deregulation of actin binding, actin filament binding and structural constitution of the cytoskeleton at the protein level with increasing severity grade. Moreover, these analyses revealed changes in severe condition for the GO term referring to tubulin-binding (Fig. 3A). To further investigate cytoskeletal changes in LIS1-patient organoids, we performed immunohistochemistry for acetylated alpha-tubulin (Ac-TUB), which is important for regulating microtubule architecture and maintaining microtubule integrity. This analysis revealed a clear overall reduction in Ac-TUB positive labeled structures in whole-tissue cleared organoids, a cellular phenotype increasing with malformation severity (Fig. 3B). We quantified this change, by investigating Ac-TUB strands in individual VZ structures in organoid cryosections. While in control organoids Ac-TUB strands were aligned from the apical to the basal side, basal strand density progressively and significantly decreased with increasing malformation severity (Fig. 3B-C, Sup. Table S11). Additionally, between patient samples, we identified a significant severity-dependent gradient in N-cadherin signal disruption, occupying a wider area from mild over moderate to severe patient-derived organoids (Fig. 3D-F, Sup. Table S12). In line with previous evidence in which we and others linked VZ-niche organization and N-cadherin signalling with ß-catenin/Wnt signalling (13) (41), GO enrichment analysis identified WNT signalling as a key pathomechanism in LIS1-lissencephaly (Sup. Fig. S3H). To monitor the onset and localization of WNT target gene activity in patient- and control-derived organoids, we developed WNT-GFP iPS cell reporter lines and differentiated them into forebrain-type organoids. We observed that the apical barrier of VZ structures from mild, moderate, and severe grade LIS1-lissencephaly patients exhibit - with increasing disease severity - a gradual decrease of WNT-target reporter activity signal compared to control (Fig. 3G-H). Changes in WNT signaling can lead to a switch of apical RG cell division from progenitor cell expansion to neurogenesis (13). Consequently, we analyzed the plane of cell division of cycling progenitors at the apical surface of control, mild, moderate, and severe grade patient-derived organoids (Fig. 3I-J). Indeed, we identified a clear increase in horizontal division patterns of apical RG cells in cultures derived from severe-grade patients compared to controls. In addition, we observed an increase in oblique division mode (most likely a direct effect of the perturbed microtubule array) in all patient samples, which appeared most prominently in patients with moderate- and severe-grade LIS1-lissencephaly (Fig. 3I). We also explored the expression of genes involved in WNT signaling in CyP, the population that is most likely to be associated with the progenitors at the apical surface, and found a major dysregulation, particularly in the severer forms of LIS1-lissencephaly (Fig. 3K).

**Fig. 3.**
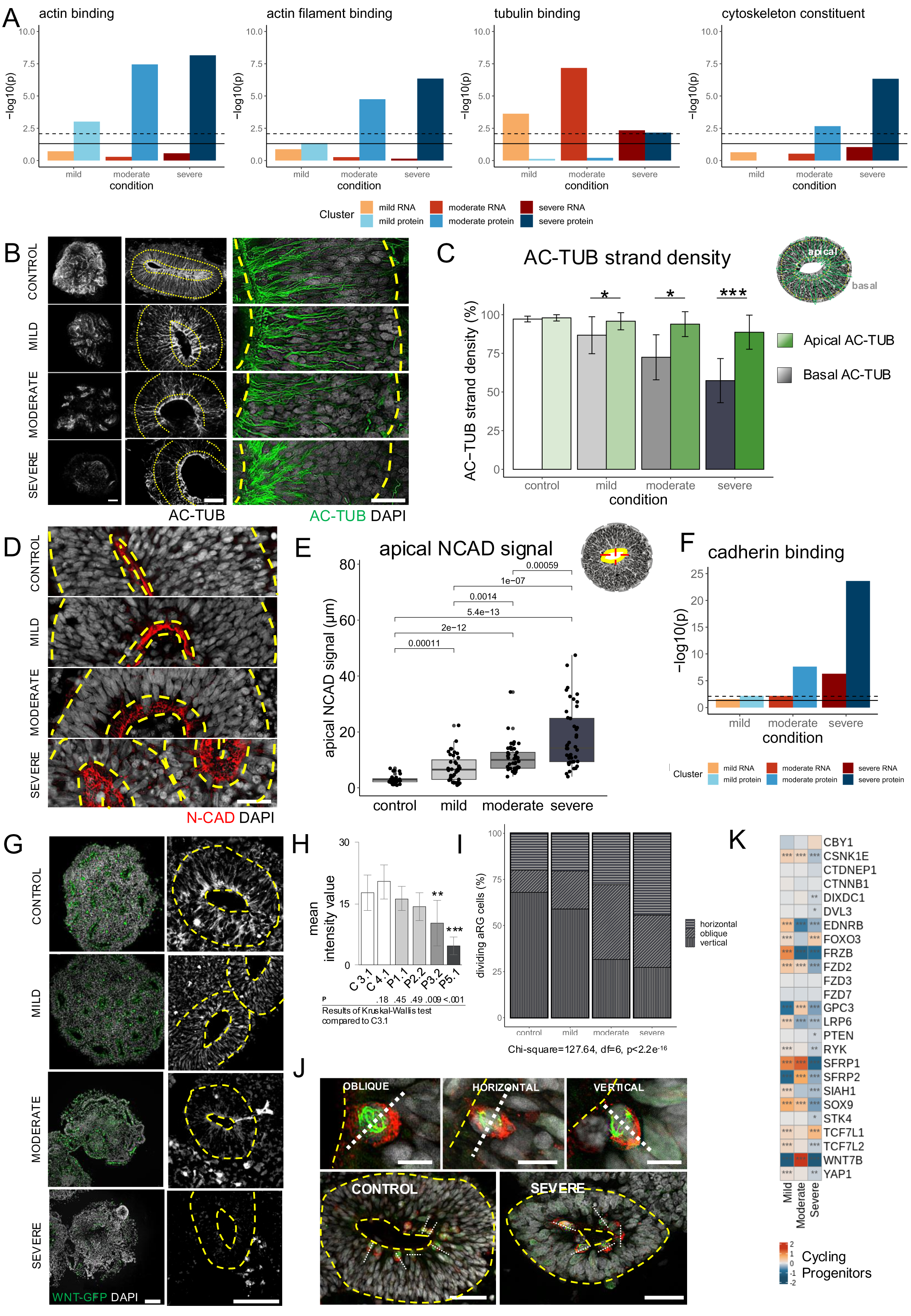
Probing selective molecular pathways identified in the OMICS analyses in LIS1-pateint-derived organoids. **A**) Bar graphs depicting the -log10-transformed p-values of the GO molecular function enrichment analysis, for the GO terms actin binding, actin filament binding, tubulin binding, and structural constituent of cytoskeleton, for differentially regulated genes (reds) and proteins (blues), solid line represents p-value threshold 0.05 and dotted line the Bonferroni-adjusted p-value threshold. **B**) Representative images of whole-tissue cleared or cryo-sectioned organoids stained for acetylated α-tubulin (Ac-TUB). Control, mild, moderate and severe LIS1-patient derived organoids at day 20. **C**) Quantification of apical and basal Ac-TUB strand density in control, mild, moderate and severe LIS1 patient-derived organoids at day 20, statistical analyses in Sup. Table S11. **D**) Representative images of control, mild, moderate and severe patient-derived organoids stained for N-cadherin (N-CAD) at day 20. **E**) Quantification of apical N-CAD signal in control, mild, moderate, and severe LIS1 patient-derived organoids at day 20, statistical analyses in Sup. Table S12. **F)** Bar graph depicting the -log10-transformed p-values of the GO enrichment analysis of the term cadherin binding, for differentially regulated genes (reds) and proteins (blues), solid line represents pvalue threshold 0.05 and dotted line the Bonferroni-adjusted p-value threshold. **G)** Representative images of WNT-GFP reporter control, mild, moderate and severe patient-derived organoids at day 20. **H**) Quantification of mean value of WNT-GFP signal in VZ structures, control N=20, mild N=20, moderate N=10, severe N=10. Error bars, ±SD. **I)** Relative quantification of cell division plane orientation at day 20, control N=14, mild N=56, moderate N=43, severe N=57 and **J)** Representative recordings of vertical-, horizontal and oblique division planes by marking dividing cells with p-VIMENTIN (green) and the mitotic spindle by TPX2 (red) in control and severe LIS1 patient-derived organoids. **K**) Heatmap showing the log2FC of differentially expressed genes belonging to WNT signaling pathway in the CyP population of each severity-grade group compared to control. *P< 0.05, **P< 0.01, ***P< 0.001. Scalebars: B: first column 200 µm, second and third column 50 µm; D: 20 µm; G: 50 µm; J: 20 µm.

### Pharmacological rescue of cellular phenotypes in the different severity grades of LIS1-patient-derived organoids

To validate key pathways perturbed in LIS1-lissencephaly organoids and investigate their relevance in mild, moderate, and severe LIS1-patient samples, we designed rescue experiments by applying the FDA-approved drug epothiloneD (a macrolide directly interacting and stabilizing microtubules) (42-44), or the GSK3ß inhibitor CHIR99021 (a WNT pathway modulator) (45). Consistently with the strong association of the severe grade LIS1 phenotype with deregulated microtubule stability, epothiloneD partially restored Ac-TUB strand density, apical N-cadherin distribution, and VZ morphological parameters, as well as reducing the neuronal belt in severe patient organoids (Fig. 4A, Sup. Fig. S4A-F, Sup. Table S13-15). These changes were less prominent in mild and moderate LIS1-patient samples and not visible in the control condition. Similarly, CHIR99021 treatment resulted in a more homogeneous generation of VZ structures, an increase in VZ diameter, and a reduced neuronal belt surrounding the VZ area, most evident in severe grade patient-derived organoids (Fig. 4B, Sup. Fig. S4G-H, Sup. Table S14). In addition, the treatment rescued the switch in the plane of cell division in organoids derived from severe-grade patients (Fig. 3I, Sup. Table S16). While CHIR99021 treatment also led to a more homogeneous generation of VZ structures in mild and moderate LIS1-patient organoids, no obvious changes could be observed in the control condition. These data support the suggested role of deregulation of WNT signaling as an underlying disease mechanism across all LIS1-patient organoids with the strongest associated with the severe LIS1-patient samples.

**Fig. 4.**
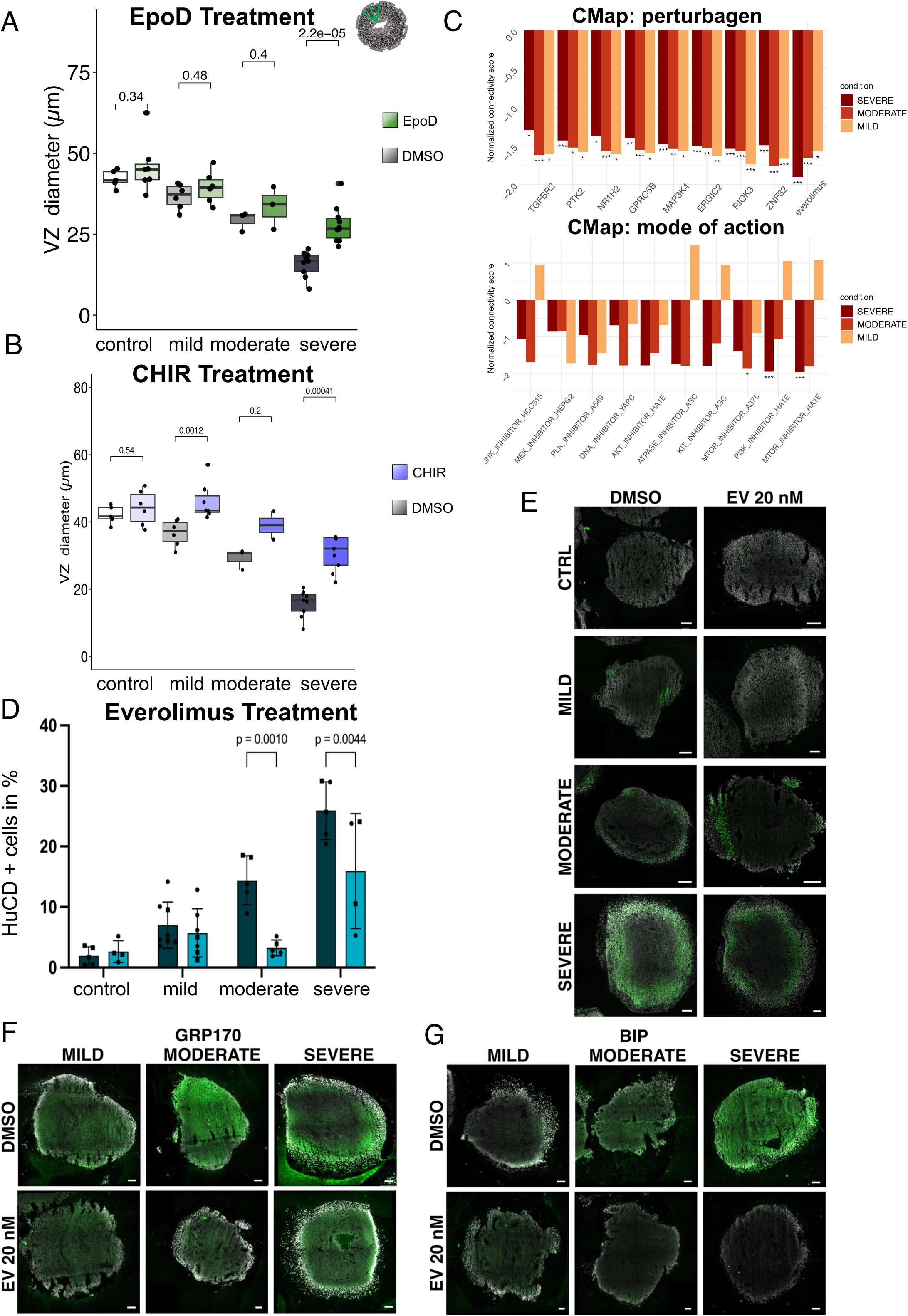
Probing therapeutic targets in LIS1 patient-derived organoids. **A**) VZ diameter quantification of DMSO and EpoD treated control- and LIS1 patient-derived organoids at day 15, statistical analyses in Sup. Table S14. Central line in boxplot represents median, lower lines the 25^th^ and upper the 75^th^ percentile, Whiskers are 1.5 interquartile ranges. Individual dots represent the mean of one batch (≥ 2 replicates). **B**) VZ diameter quantification of DMSO and CHIR treated organoids. Central line in boxplot represents median, lower lines the 25^th^ and upper the 75^th^ percentile, Whiskers are 1.5 interquartile ranges. Individual dots represent the mean of one batch (≥ 2 replicates). **C)** Barplots depicting connectivity scores from CMap analyses of perturbagen classes (left) and modes of actions (right), based on the differentially expressed genes between the mild, moderate, severe and control conditions, *p< 0.05, **p< 0.01, ***p< 0.001. **D)** HuCD positive cells quantification of DMSO and everolimus treated control and LIS1 patient-derived organoids at day 20; data are presented as mean ± SD. **E)** Representative images of organoids of different severities stained for HuCD upon everolimus treatment. **F**) Representative images of organoids of different severities stained for GRP170 upon everolimus treatment **G**) Representative images of organoids of different severities stained for BiP upon everolimus treatment. Scale bars: 125 µm.

To identify rescue strategies for the phenotypic changes in LIS1-patient organoids identified in the progenitor cell omics data, we analyzed the gene expression signature of the RG cluster in LIS1-patient derived organoids against a library of compound-induced profiles using the Connectivity Map (CMap) approach. This analysis identified mTOR inhibition as the most significant mode of action with a negative connectivity score (Fig. 4C), highlighting everolimus as the perturbagen with the greatest potential to rescue LIS1-lissencephaly-associated transcriptional alterations in LIS1-patient organoids. Everolimus has an analogous function to that of rapamycin that, by inhibiting mTORC1, reduces the production of downstream effectors S6K1 and 4E-BP1 and reduces protein synthesis (46). It has been approved by both the EMA and FDA as treatment for multiple cancers and, more recently as an add-on treatment in patients with seizures related to tuberous sclerosis complex that have not responded to other treatments (47, 48). In this context, data indicates that it can reduce abnormal brain cell growth and over-proliferation, correct irregular neuronal spatial organization, enhance neuronal myelination, mitigate abnormalities in neuronal morphology, improve synaptic plasticity, and decrease excessive inflammatory mediators in the cerebral cortex (49). Treatment of LIS1-patient organoids with everolimus led to a significant reduction in HuC/D positive cells (a pan-neuronal marker), with a 4.5-fold decrease in moderate conditions and a 1.6-fold decrease in severe conditions, compared to the respective DMSO-treated samples suggesting a reduction in premature neurogenesis (Fig. 4D-E). Notably, expression of the ER chaperone BiP (HSPA5) or its nucleotide exchange factor GRP170 (HYOU1), both also strongly expressed in the moderate and severe cases, were also reduced upon everolimus treatment (Fig. 4F-G). Thus, our data suggest for the first time that mTOR inhibitors represent interesting candidates to develop novel therapeutic strategies for moderate and severe grade LIS1-lissencephaly in humans.

## Discussion

In the current study, we use LIS1-lissencephaly as a model to address a major challenge for cerebral organoid research: the ability to capture disease severity, an important step towards implementing organoids in personalized medicine and drug discovery. LIS1-lissencephaly is particularly suitable to probe this question since it is caused by a mutation in the *LIS1* gene, it presents with different severity grades, it is a malformation associated with structural changes of human brain tissue, suitable animal models remain complicated and access to affected primary tissue at early developmental time points is not feasible.

Our study demonstrates that it is possible to recapitulate the development of disease severity *in vitro* and that the grade of phenotypic changes aligns with the clinical severity grade of the patients. Thus, we suggest that our observations are driven primarily by the severity grade rather than background genetic differences. Further, we found that some changes in molecular pathways are restricted to patients with moderate and/or severe LIS1-lissencephaly. In contrast, we also found that LIS1-patient-cells display independent of the severity grade premature neurogenesis and proteotoxic stress illustrated by depletion of NE cells and upregulation of protein catabolism, respectively. Of note, the proliferative capacity and the neurogenic potential of NE and RG cells are among the cardinal pillars of the expansion and development of the human gyrencephalic brain (50), which is strongly perturbed in all LIS1-lissencephaly patients. Further, we note that the increase in severity grade strongly correlates with an increase in microtubule network alterations. We were able to show a reduced expression level of the LIS1 protein in each condition, confirming that a LIS1 dosage effect may contribute to microtubule network alterations. Importantly, cytoskeletal destabilization and disorganization of the tubulin network were also evident and misregulation of TUBB, TUBG1, and dynein complex members were previously identified (51-53) which we could confirm related to progressive severity of the phenotype in LIS1 patient cells. Given the role of microtubules in N-cadherin recycling (54), the degree of its disruption likely impacts the degree of the impairment of the adherens junction scaffold, leading to VZ-niche dis-organization which can converge into changes in WNT-signaling (13, 55) a pathway essential for neurogenesis and progenitor expansion (56) and confirmed to be significantly perturbed in severe LIS1-patient organoids.

Disrupted cytoskeletal arrangement has been also linked to the build-up of protein aggregates based on impaired vesicular transport, in turn resulting in the activation of different cellular stress response mechanisms such as UPR and proteolysis (57). Activation of UPR in mice was described to induce direct neurogenesis resulting in microcephaly (58), but never linked to LIS1-lissencephaly. Notably, several proteins involved in alternative splicing, mRNA stability, and translation, such as DDX3X, NCL, SRSF1, SSB, and FUS were significantly upregulated in severe LIS1-patient samples. Further, proteomic data unveiled significant alterations in proteostasis in the severe grade LIS1-patient organoids, marked by an increase of proteins involved in proteolysis and unfolded protein binding, including BiP (the major chaperone resident in the ER) and its co-factor GRP170/HYOU1 as well as HSP90-alpha, HSP90-beta and HSP90B1 (endoplasmin/GRP94), along with the HSP90 co-chaperones CDC37 and DNAJC7. Additionally, the lectin chaperones calnexin and calreticulin, along with various disulfide isomerases and DNAJC8, were upregulated suggesting an increased folding activity in the ER and the activation of protective mechanisms similar to those seen in neurodegenerative diseases such as Parkinson’s and Machado-Joseph disease (59-62). Notably, our treatment with everolimus was able to correct the elevated levels of BiP and GRP170, suggesting that mTOR modulation can restore aspects of proteostasis dysregulation in severe LIS1-patient organoids.

As well as being able to validate the role of cytoskeletal instability and Wnt/ß-catenin signaling in severe LIS1-patient-derived organoids, we for the first time associate moderate and severe LIS1-lissencephaly with perturbances in the mTOR signaling cascade by rescue experiments, applying ephotiloneD, CHIR99021, or everolimus, respectively. Interestingly, Wnt/ β-catenin pathway inhibition has been associated with the activation of the PI3K/AKT/mTORC1 pathway (63), which we could confirm by our drug repurposing analysis. Such conditions may impair WNT-dependent progenitor proliferation, favoring premature neuronal differentiation while depleting the progenitor pool, explaining the severity-dependent alterations in progenitor cell homeostasis and neurogenesis observed in LIS1-organoids. CHIR99021 treatment, by inhibiting GSK3β, activates WNT signaling through β-catenin accumulation and ameliorates the loop parameters. In contrast, Everolimus inhibits mTORC1 through FKBP12, leading to MNK-mediated phosphorylation of eIF4E (64), allowing β-catenin translocation into the nucleus (65) independently of upstream Wnt signaling. As a result, everolimus could relieve proteotoxic stress through mTOR modulation, while partially alleviating WNT impairment, resulting in reduced differentiation.

MCDs leading to treatment-refractory epilepsies have already been associated with dysregulation of Wnt/ß-catenin and mTOR signaling. Specifically, focal cortical dysplasia type II (FCDII), tuberous sclerosis complex (TSC), and hemimegalencephaly (HME), and Rett syndrome reveal a disease-specific association of increased mTOR activity (66, 67). Inversely, very recently, mutations in the p53-induced death domain protein 1 (*PIDD1*) and the contiguous gene Miller Dieker syndrome (MDS) were associated with a decrease of mTOR signaling (15), indicating that deregulation of mTOR in general can lead to similar spectrums of brain malformation and symptoms. Further, MDS and hippocampal sclerosis (HS) have been associated by us and others with dysregulation of Wnt/ß-catenin signaling (68). In line with this, both pathways have been suggested as novel therapeutic targets aiming to either block or reduce seizures or modulate the underlying course of the disease and, by that, prevent symptoms (15, 68). Considering that the different MCDs have been connected to deregulation in mTOR and Wnt/ß-catenin one might speculate that MCDs with different pathophysiology may share common underlying principles and that perturbance in the mTOR pathway might be a major contributor to pathological dysfunction. Along with this hypothesis, studies investigating FCDII, TSC, and HME as well as our study on LIS1-lissencephaly, exemplify mTOR inhibitors as a potential new therapeutic strategy for a heterogeneous group of focal or diffuse developmental brain malformations leading to childhood-to adulthood-onset pharmaco-resistant epilepsies (66, 69). Despite this, it needs to be examined in future studies whether here identified pathways involved in the development of the different LIS1-lissencephaly severity grades, persist in progenitor derivates i.e. in neuron and glial cells relevant for the patients’ post-embryonic development as observed for FCDII, TSC, HME, and HS. Furthermore, it will be interesting to validate the additional drugs identified in our drug repurposing analysis including those suggested to be more selectively effective for either mild, moderate, or severe LIS1-lissencephaly.

Finally, our results may advance efforts to model the biology of severity in other brain disorders with much larger cohorts of patients-derived organoids for a given disease, without “*a priori*” clinical information on disease severity and conducted under fully blinded conditions. These studies can confirm molecular principles or identify new signaling pathways that have not yet been linked to the given disease. Thus, this model may also confirm patient stratification or provide additional information leading to new mutation-specific subgroups not previously envisioned in the clinical dataset, and thus could boost the development of more effective therapies.

## Acknowledgement

We thank Isabell Moskal, Gina Tillmann, Helene Schamber and Elina Nürnberg for the pivotal technical support. We thank the DKFZ Single-Cell Open Lab (scOpenLab) for assistance with the scRNA sequencing experiment. We acknowledge the support of the NGS Core Facility Mannheim, Medical Faculty Mannheim of Heidelberg University.

## Declarations

### Author Contributions

Conceptualization, L.Z., A.C.R., O.F., S.G.M., P.K., F.F., and J.L.; Methodology, L.Z., A.C.R., O.F., M.G., C.M., A.Ho., E.Z., A.J., F.M., R.W., C.B.S., A.He., A.A, S.H.W, T.R., A.R., N.B-B. and J.L.; Validation, L.Z., A.C.R., O.F., M.G.; Formal Analysis, L.Z., O.F., M.G.; Investigation, L.Z., A.C.R., O.F.; ScRNA-Seq Data analysis, L.Z., E.Z. A.Ho., A.C.R., J.L.; mass spectrometry including data analyses C.B.S., A.He., T.R., A.R., M.G., L.Z., J.L., Whole-exome-sequencing, C.M. and N.B-B.; Writing – Original Draft, O.F., J.L.; Writing – Reviewing & Editing, L.Z., A.C.R., M.G., A.R., N.M., S.G.M., F.F. and J.L.; Visualization, L.Z., A.C.R., O.F. M.G, J.L.; Supervision, J.L.; Project Administration, J.L.; Funding Acquisition, A.He, N.B-B., F.F., J.L.

### Data and code availability

Raw data will be made available through the European Genome-Phenome Archive. All original code has been deposited at GitHub repository and is publicly available as of the date of publication (https://github.com/lzillich/LIS1_code).

### Inclusion and Ethics

For the human samples used in the study, all participants provided written informed consent prior to participation in the study. This study was ethically approved by the committees of the Imagine - Institut des maladies génétiques, Paris (approval number no. DC2014-2272) and by the Ethics Committee II of Medical Faculty Mannheim of Heidelberg University (approval no. 2014-626N-MA).

## Materials and methods

### Cell lines and patients

LIS1-patient fibroblasts and lymphocytes were collected from Nadia Bahi-Buisson from the Necker Enfants Malades University Hospital in France. We selected those LIS1 patients in this study for which blood and/or fibroblast samples were available, which were clinically and genetically well-characterized, for which high-quality MRI data were available, and for whom patient consent had been obtained. For more details see Philbert, Maillard (19) (LIS1-mild 1, 8-year-old male donor, c.569-10T>C LIS1 mutation; LIS1-mild 2, 5-year-old male donor, c.569-10T>C LIS1 mutation; LIS1-moderate 1, 6-year-old female donor, c.13delC LIS1 mutation; LIS1-moderate 2, 13-year-old female donor, delEx11 LIS1 mutation; LIS1-severe 1, 4-year-old female donor, c.1002+1G>T; LIS1-severe 2, 18-year-old female donor, c.531G>C LIS1-mutation; LIS1-severe 3, 3-yea-old female donor, c.445C>T LIS1 mutation). Each patient-specific LIS1 mutation was confirmed using Sanger sequencing after PCR amplification of the coding sequences. 2 control iPS cells were derived from skin fibroblasts obtained from Coriell Biorepository (control 1, 2-years old female, catalog ID GM00969, control 2, 5-month-old male, catalog ID GM08680) and addition, 5 control iPS cell lines were received from Dr. Sandra Horschitz (Ethics Committee II of Medical Faculty Mannheim of Heidelberg University approval no. 2014-626N-MA, control 3, 21-year-old female donor; control 4, 44-year-old female donor; control 5, 25-year-old female donor; control 6, 26-year-old female donor, control 7, 23-year-old male donor). It is important to note that there are no known sex-specific defects associated with the conditions being studied, and age-related cellular features are thought to be largely reset after reprogramming (70).

A complete overview of the cell lines used in this study is provided in Supplementary Text S1.

### Reprogramming of fibroblasts and lymphocytes

Somatic cells were reprogrammed by non-integrative delivery of OCT4, SOX2, KLF4, and c-MYC using the CTS^TM^ CytoTune^TM^-iPS 2.1 Sendai Reprogramming Kit (Thermo Fisher). The manufacturer’s instructions were strictly followed (CTS^TM^ CytoTune^TM^-iPS 2.1 Sendai Reprogramming Kit User Guide).

### iPS cell validation and culture

Pluripotency of reprogrammed cells was quality-controlled by detection of the pluripotency-associated markers TRA-1-60, TRA-1-81, and SSEA-4 by immunocytochemistry and their capacity for spontaneous differentiation into cell types of all three germ layers. To induce 3-germ-layer differentiation iPSCs were dissociated into single cells using TrypLE Express and plated in an ultra-low-binding 96-well plate (9000 cells/well; Amsbio, lipidure-coat plate A-U96) in Pluripro-medium (PP, Cell guidance systems) supplemented with 50 µM ROCK inhibitor to induce embryoid body (EB) formation. After 2 days, Ebs were plated onto extracellular matrix (GT)-coated dishes in DMEM containing 10% fetal calf serum (FCS), 1% pyruvate, and 1% non-essential amino acids (NEAA, Gibco). Cells were cultured for 4 weeks before being subjected to immunocytochemical analysis. Further whole-genome single nucleotide polymorphism (SNP) genotyping was performed for each iPS cell line for karyotyping. To that end, genomic DNA was prepared using the Dneasy Blood & Tissue Kit (Qiagen). SNP genotyping was performed at the Institute of Human Genetics at the University of Bonn. Genomic DNA at a concentration of 50 ng/µl was used for whole-genome amplification. Afterward, the amplified DNA was fragmented and hybridized to sequence-specific oligomers bound to beads on an Illumina OmniExpressExome v1.2 chip or Illumina Infinium PsychArray-24 v1.1 chip. Data were analyzed using Illumina GenomeStudio V2011.1 (Illumina). Patient-specific LIS1 mutations were validated in the iPS cells by Sanger sequencing for the LIS1-mutation c.13delC and c.569-10T>C, and in fibroblasts by Multiplex Ligation-dependent Probe Amplification (MLPA) for the LIS1-mutation: exon 11 del, c.1002+1G>T and c.445 C>T. Induced PS cells were maintained on Geltrex-coated cell culture plates (Thermo Fisher) in PP medium or Essential 8 (E8) medium at 37 °C, 5% CO2, and ambient oxygen level with daily medium change. For passaging cells were treated with TrypLE Express (Stem Cell Technologies) or EDTA (Thermo Fisher Scientific). After passaging, medium was supplemented with 5μM Y-27632 (CellGuidance Systems) to foster cell survival. All human iPS cell lines were regularly tested and confirmed negative for mycoplasma.

### Generation of WNT-GFP reporter iPS cell lines

iPS cells were transduced with a Lentivirus expressing GFP under activation of WNT signaling (Lentiviral-Top-dGFP reporter, Addgene plasmid #14715). Puromycin (1μg/ml, Sigma-Aldrich) selection was initiated 48 h following transduction. iPS cell-WNT reporter lines were used for forebrain-type organoid generation.

### Generation of 3D forebrain-type organoids and cortical progenitor cells

Cerebral forebrain-type organoids were generated and quality controlled as previously described (20, 25). Briefly, iPS cell colonies were dissociated using TrypLE Express (Thermo Fisher Scientific) and cells were plated in u-bottom 96-well plates previously coated with 5% Pluronic F-127 (Sigma Aldrich) in phosphate-buffered saline (PBS). Cells were seeded in a volume of 150μl/well in PP or E8 medium supplemented with 50μM Y-27632. Following successful cell aggregation, medium was switched at day 5 to neural induction medium (DMEM/F12, B27 supplement 1%, N2 supplement 0,5%v/v, cAMP 300ng/ml, GlutaMAX 1%, NEEA 1%, D-Glucose, Insulin, KOSR 2%, Penicillin/Streptomycin 1% v/v, heparin 1µg/ml, LDN-193189 0,2mM, A83-01 0,5mM and XAV 2µM) with a medium change every second day. Sufficient neural induction was monitored by the development of translucent and smooth edges using a bright field microscope. On days 9–11, when translucent neural ectoderm was visible, organoids were embedded in GT (Thermo Fisher), transferred to Pluronic F-127-coated dishes and maintained in neural differentiation medium (DMEM/F12, B27 supplement 1%, N2 supplement 0,5% v/v, cAMP 300ng/ml, GlutaMAX 1%, NEEA 1%, D-Glucose, Insulin, KOSR 2% and Penicillin Streptomycin 1% v/v). After this passage, organoids were kept in 10cm or 6cm dishes under agitation at 70rpm on an orbital shaker (Infors Celltron HD) at 37°C, 5% CO_2,_ and ambient oxygen level. Media was changed every 3 to 4 days. At day 15 or 20, organoids were harvested for phenotypical analysis, while for scRNA-seq they were collected at day 20+/-3. For immunostaining, 20µm sections were prepared using a cryotome. At least six organoids for each of three different batches were analyzed.

Induced PS cell-derived neural progenitors were generated along established protocols (25) with slide adaptations. Neural induction was initiated in 90-95% confluent iPS cell cultures by changing the culture medium to neural base medium containing advanced DMEM/F12 (Thermo Fisher Scientific, 12634010) supplemented with 1% Pen/Strep, Glutamax (1x; Thermo Fisher Scientific, 35050038) and 1% B-27 supplement (Thermo Fisher Scientific; 17504044). Cells were grown in neural base medium supplemented with SB-431542 (10 µM; Cell Guidance Systems, SM33), LDN-193189 (1 µM; StemCell Technologies, 72148) and XAV939 (2 µM; Cell Guidance Systems, SM38) for 8 days. On day 8 (in case of the LIS1 moderate line 1.2. on day 6 as this line tend to premature differentiate) neural progenitor base medium was supplemented with LDN-193189 (200 nM) and XAV939 (2 µM). Cells were harvested for mass spectrometry 2 days after passaging and replating in neural differentiation media containing DMEM/F12, 0.5% N2 supplement, 1% B27 supplement, and cAMP (300 ng/ml) on GT-coated cell culture plates. Cells cultured in parallel were used for quality control which include immunohistochemistry for neural progenitors and neurons. Only those batches where homogenous neural induction was confirmed were further processed for mass spectrometry. At least 4 samples derived from at least two different genetic backgrounds and generated from at least three different batches were processed for mass spectrometry.

### CHIR, epothiloneD and everolimus Treatment

Organoids were treated after the induction phase, from culture day 10 to day 15 with 1µM CHIR or 1nM epothiloneD. For treatment with everolimus, organoids were treated from day 10 to day 19 with 20 nM everolimus, refreshing media every three days and fixed at day 20. The drugs were resuspended in DMSO to reach the concentration of 1µM CHIR, 1nM epothiloneD or 20 nM everolimus, respectively. After treatment, organoids were fixed and cryo-sectioned for immunocytochemical analysis.

### Clearing of 3D forebrain-type organoids

For whole tissue mounting, organoids were fixed with 4% PFA for 2 h at RT and optically cleared according to Susaki et al. (71). Blocking was done with 10% horse serum, 0.2% gelatin from cold water fish skin, and 0.1% Triton X-100 diluted in PBS for 24 h at 37°C, followed by primary antibody incubation for 48 h at 37°C. Secondary antibody incubation was done for 48 h at 37°C. Refractive index matching was performed according to Nürnberg et al. (72) by immersion of samples in an aqueous solution of glycerol (RI=1.457) for 48 h at RT. Samples were mounted in U-shaped 2.5 mm glass capillaries by embedding in 0.1% low melting agarose in ddH_2_O. For light sheet microscopy, glass capillaries were transferred into 35 mm glass bottom dishes, immobilized by agarose embedding, and immersed in RI-matched glycerol solution. For temperature adjustment, samples were kept in the microscopy room for at least 24 hours before image acquisition. Image acquisition was done using a Leica Microsystems TCS SP8 DLS, equipped with LAS X software, L 1.6x/0.05 DLS illumination objective, HC APO L 10x/0.30 W DLS detection objective and 7.8 mm Glycerol DLS TwinFlect mirrors. Image stacks were acquired with a step size of 3.7 µm and fused with LAS X.

### Histology and Immunocytochemistry

Cells and organoids were fixed with 4% paraformaldehyde (PFA) for 10 min at room temperature (RT) and blocked in 10% Fetal Calf Serum in PBS with 0,1% Triton for 1 h at RT. Primary antibodies were diluted according to the manufacturer’s instructions and incubated overnight at 4°C with the following dilutions: Ac-TUB (1:500, Cell signaling), AFP (1:200, Hölzel), TUBB3 (1:2000, Cell Signaling), NANOG (1:200, DSHB), N-CAD (1:500, BD), p-VIM (1:500, Novus biologicals), TPX2 (1:500, Novus Biologicals), OCT3/4 (1:500, R&D Systems), SMA (1:400, Abcam), SSEA3 (1:500, Abcam) and SOX2 (1:500, Santa Cruz), HSP90 (1:250, Genetex), BiP (1:250, provided by Professor Martin Jung, University of Saarland). The secondary antibodies were diluted according to the manufacturer’s instructions and incubated for 1h at RT (488-ms/rb; 555-ms/rt; 647-ms, 1:1000, Invitrogen). Nuclei were visualized using 0,1μg/ml DAPI (Sigma Aldrich). Stained sections were stored at 4°C and imaged using the Inverted Leica DMIL LED Microscope with the Thunder imaging software (Leica).

### Organoid quantifications

Organoid quantifications were performed as previously described (13) with slight adaptations. Images were acquired using the Inverted Leica DMIL LED Microscope with the Thunder imaging software (Leica) and analyzed using ImageJ software. All quantifications were done in at least three organoids with at least six VZ-like structures for each of at least three different organoid batches. The given n number is the total of VZ-like structures analyzed. For the quantification of the VZ dimension parameters, sections were stained with DAPI. Length and area measurements were performed with Image J software. For the VZ diameter, three measurements for each cortical VZ structure were performed: one straight measurement from the apical to the basal side of the VZ structure and two measurements both starting at the apical point of the first measurement while the basal point was located in a 45-degree angle on the right or left side. The mean of the three values was taken as VZ diameter. In the case of heterogenous VZ diameter, the thickest area was considered. The VZ tissue area was defined as the ratio of the total VZ area to the ventricle area. Ac-TUB strand density was measured by plot profile determination using ImageJ software and a self-designed Excel file containing formulae for background subtraction and automatic signal peak counting. The disruption diameter of N-CAD was measured at four apical membrane positions (90, 180, 270, and 360 degrees) using ImageJ software. The mean value was taken as disruption diameter. Mitotic spindles were analyzed by immunocytochemical staining using p-VIM for marking dividing RG cells at the apical membrane and TPX2 for the visualization of the mitotic spindle.

### Cortical organoid dissociation for single-cell RNA-sequencing

20 +/- 3-day old organoids (two or three per condition: control C1, control C7, LIS1-mild P1, LIS1-mild P2, LIS1-moderate P3, LIS1-moderate P4 and LIS1 severe P5, LIS1-severe P6) were sliced with a scalpel and dissociated according to our already published protocol (25). Briefly, the tissue was incubated in papain (Sigma Aldrich) containing buffer (1mM L-cysteine and 0,5mM EDTA in Earle’s balanced salt solution, 20 units of papain, and 10μg/ml of DNase (Sigma Aldrich) for 20 min at 37°C. After incubation, organoids were washed with differentiation media and dissociated mechanically using a 1% bovine serum albumin (BSA)-coated 1000µl pipette. After centrifugation at 400g for 4 min at 4°C, the cell pellet was resuspended in 1ml ice-cold PBS supplemented with 0.04% BSA and filtered through a 30μm cell strainer. Single cell library preparation was performed using the 10x Genomics Chromium platform according to the 10x Genomics Chromium Single Cell 3’ Library & Gel Bead Kit v3.1 chemistry user guide (10x Genomics). The prepared cDNA libraries were processed by the High Throughput Sequencing Unit of the Genomics & Proteomics Core Facility of the German Cancer Research Centre (DKFZ). The libraries were sequenced on two lanes of the Illumina NovaSeq 6k platform on a S1 flow cell (paired-end 28+94 bp).

### ScRNA-seq data quality control and preprocessing

Count matrices for single-cell RNAseq data were generated from fastq files using cellranger (10x Genomics). Data analysis was performed using Seurat v.4.0.5 (73), if not stated otherwise. During quality control, features that were not expressed in any cell were removed from the count matrix. Next, cells were removed in each replicate individually based on the number of expressed features, total UMI counts, and mitochondrial gene fraction (>10%). For detailed QC parameters, see Sup. Table S17. For each sample, normalization was performed using *sctransform*. Cell cycle differences (S.Score-G2M.Score) were regressed out and cell multiplets were removed using the R package DoubletFinder v.2.0.3 (74). Integration of data was performed in two steps: first, for every condition, the two samples were integrated, followed by an integration of the four integrated objects resulting in the final Seurat object. UMAP dimensionality reduction and nearest-neighbor graph construction were performed based on dims 1:50. A resolution threshold of 0.15 was used for cluster generation. Cluster identity was determined based on the expression of known cell type markers. The intermediate progenitor (IP) cluster was manually split into two clusters according to normalized expression levels of the EOMES gene (>0.25) yielding a total of 10 clusters in the final object.

### Sample preparation for mass spectrometry

Proteins from cortical progenitors were extracted from TriFast homogenized samples following manufacturer’s protocol. Dried, snap-frozen protein pellets, underwent lysis through the addition of 200 µl of a 50 mM Tris-HCl buffer (pH 7.8) containing 5% SDS and cOmplete ULTRA protease inhibitor (Roche). The samples were then subjected to Bioruptor® treatment (Diagenode) for 10 minutes (30 sec on, 30 sec off, 10 cycles) at 4 °C, followed by centrifugation at 4 °C and 20,000 g for 15 minutes. Subsequently, the protein concentration in the supernatant was determined using the BCA assay according to the manufacturer’s guidelines. Disulfide bonds were reduced with the addition of 10 mM TCEP at 37 °C for 30 minutes, and free sulfhydryl bonds were alkylated with 15 mM IAA at room temperature in the dark for 30 minutes. For proteolysis, 100 µg of protein from each sample was utilized following the S-trap protocol (Protifi). The protein-to-trypsin ratio was maintained at 20:1, and digestion took place for 2 hours at 45 °C. The proteolysis process was terminated by acidifying the sample with formic acid (FA) to achieve a pH below 3.0. Verification of complete digestion of all proteolytic digests occurred post-desalting, utilizing monolithic column separation (PepSwift monolithic PS-DVB PL-CAP200-PM, Dionex) on an inert Ultimate 3000 HPLC (Dionex, Germering, Germany) through direct injection of 1 μg of the sample. A binary gradient (solvent A: 0.1% TFA, solvent B: 0.08% TFA, 84% ACN) ranging from 5-12% B in 5 minutes and then from 12-50% B in 15 minutes at a flow rate of 2.2 μL/min and at 60 °C was applied, with UV traces recorded at 214 nm (75).

### Mass Spectrometry sample analysis

A total of 1 g of the respective peptide samples underwent separation using an Ultimate 3000 Rapid Separation Liquid Chromatography (RSLC) nano system equipped with a ProFlow flow control device, in conjunction with a Q Exactive HF Orbitrap mass spectrometer (Thermo Scientific, Schwerte, Germany). Peptide concentration employed a trapping column (Acclaim C18 PepMap100, 100 μm, 2 cm, Thermo Fisher Scientific, Schwerte, Germany) with 0.1% trifluoroacetic acid (TFA) from Sigma-Aldrich, Hamburg, Germany, at a flow rate of 10 L/min. Subsequent reversed-phase chromatography (Acclaim C18 PepMap100, 75 µm, 50 cm) utilized a binary gradient (solvent A: 0.1% formic acid (Sigma-Aldrich, Hamburg, Germany); solvent B: 84% acetonitrile (Sigma-Aldrich, Hamburg, Germany) with 0.1% formic acid; 5% B for 3 min, linear increase to 25% for 102 min, a further linear increase to 33% for 10 min, and a final linear increase to 95% for 2 min followed by a linear decrease to 5% for 5 min). For MS survey scans, the parameters included operating MS in data-dependent acquisition mode (DDA) with full MS scans from 300 to 1600 m/z (resolution 60,000) and the polysiloxane ion at 371.10124 m/z as a lock mass. The maximum injection time was set to 120 ms, and the automatic gain control (AGC) was set to 1E6. Fragmentation involved selecting the 15 most intense ions (above the threshold ion count of 5E3) at a normalized collision energy (nCE) of 27% in each cycle, following each survey scan. Fragment ions were acquired (resolution 15,000) with an AGC of 5E4 and a maximum injection time of 50 ms. Dynamic exclusion was set to 15 s.

### Mass spectrometry data preprocessing

All MS raw data underwent processing using Proteome Discoverer software version 2.5.0.400 (Thermo Scientific, Bremen, Germany) and were subjected to a target/decoy mode search against a human Uniprot database (www.uniprot.org, downloaded on 21 November 2019) utilizing the MASCOT and Sequest algorithm. The search parameters included precursor and fragment ion tolerances of 10 ppm and 0.5 Da for MS and MS/MS, respectively. Trypsin was designated as the enzyme with a maximum of two allowed missed cleavages. Carbamidomethylation of cysteine was set as a fixed modification, and oxidation of methionine was set as a dynamic modification. Percolator false discovery rate (strict) was established at 0.01 for both peptide and protein identification. A Label-free Quantification (LFQ) analysis was conducted for proteins with a minimum of two unique peptides, encompassing replicates for each condition.

### Statistical analysis

All statistical tests were performed two-sided with an alpha-error of 0.05 to indicate nominal statistical significance.

All quantitative data generated on organoids following immunocytochemistry was performed in batches of at least duplicates, at least two batches per cell line were analyzed. The number of included individual data points and batches can be found in the tables summarizing each statistical analysis. On average, we observed 6.26 individual data points per batch for VZ-like parameters, 4.07 for Ac-TUB, 3.67 for N-CAD disruption diameter, and 3 for cell division quantifications. During one routine karyotyping, we discovered abnormalities in cell line C2. Therefore, this cell line was excluded from all statistical analyses. Raw data was aggregated per batch, after which conditions were compared using Kruskal-Wallis- and *post-hoc*-pairwise Wilcoxon tests, as all observed parameters did not follow a gaussian distribution, which was tested using the Kolmogorov-Smirnov-Test. Differences in proportions were tested using Chi-Square tests. All results were corrected for multiple testing using the Bonferroni correction, if not stated otherwise.

For differential expression testing in the single cell sequencing data, an FDR-corrected p-value of 0.05 and an absolute log2FC larger than 0.5 was used to indicate statistical significance. Differential gene expression between the severities and the control condition in the progenitor cell populations (RG, NE, t-RG) was tested using the *FindMarkers()* function in Seurat, which encompasses a Wilcoxon Rank sum test. The minimum percentage of gene expression was set to 0.25, meaning that a gene had to be expressed in at least 25% of cells in each population, therefore avoiding false positives by not testing genes with low expression.

Protein abundances were normalized, and log-transformed. To compare the severity grades with the control condition, Wilcoxon Rank sum tests were performed. Effect sizes were calculated by dividing the Z statistic through the square root of the sample size. FDR-correction was applied to correct for multiple testing and results were filtered for an absolute effect size larger than 0.75. In addition to testing differential protein regulation, we performed a weighted correlation network analysis using WGCNA (version 1.72-1) (76). Here, networks of co-regulated proteins are constructed based on the covariance of the normalized protein abundances. We used a soft threshold of 8, a signed TOM matrix, a minimum module size of 30 and a maximum block size of 20,000 to construct the modules. In WGCNA, modules are assigned random colors. Module eigengenes were correlated with the disease severities compared to the control condition. For significantly associated modules, in at least one of the conditions, a GO enrichment analysis was performed using the approach described below. For module hubgenes of the LIS1-associated modules, we constructed protein-protein interaction networks using the STRING database (v.11.5) (77).

Integrative Gene Ontology enrichment analysis of cellular pathways (GO) Biological Processes and Molecular Functions) of differentially regulated genes (p_adj_<0.05) and proteins (p<0.05) was determined using the enrichGO() and compareCluster() functions of the R package clusterProfiler (v. 4.6.2). For this purpose, genes contributing to a GO term were retrieved using the org.Hs.eg.db package (v.3.14.0). We determined semantic overlap of enriched GO-terms using the pairwise_termsim() function of enrichplot (v.1.18.3) and visualized the results using the emapplot() function of the same package.

### Drug repositioning analysis

The NIH LINCS L1000 database (78) was used as a reference dataset for drug repositioning analysis in connectivity map analysis (CMap, https://clue.io, software version 1.1.1.43). The top 150 upregulated and downregulated DE genes from the RG cluster were used to generate the maximum input size for the CMap query tool. CMap allows an *in-silico* drug repositioning analysis by comparing expression changes of n=978 landmark transcripts in response to standardized drug treatment protocols against the query differential expression signature. Resulting connectivity scores indicate rescue (negative score) or aggravation (positive score) of queried transcriptional changes by drug treatment. Next to the generation of connectivity scores for individual drugs or more general *“perturbagens”*, CMAP also provides connectivity information at the perturbagen class, mechanism of action, and pathway level thus enabling a more general understanding of potential therapeutic targets in a phenotype. For visualization of CMap results, waterfall plots were generated using ggplot2 (v.3.4.2). Results for individual perturbagens such as pharmaceutical drugs or gene products were filtered based on negative normalized connectivity scores (NCS<0) in all three conditions (mild, moderate, severe). Mechanism of action (MOA) results were filtered based on negative normalized connectivity scores in at least two conditions (moderate and severe).

---

**Supplementary Figure 1: LIS1-patient-cohort and iPS cell generation. A**) Table containing patient information including age, sex, LIS1 mutation and severity grade. **B-E**) Representative pictures of iPS cell from LIS1-patient P2 clone 1 stained for SOX2, OCT3/4, NANOG and SSEA3. **F-G**) Representative immunocytochemical images of 4-weeks spontaneously differentiated iPS cells derived from LIS1-patient P2 clone 1 stained for the mesoderm marker smooth muscle actin (SMA) and the ectoderm marker beta-III-tubulin (TUBB3). **H**) High resolution single nucleotide polymorphism analysis was performed to control chromosomal integrity after reprogramming. For every chromosome of LIS1-patient P2 clone 1 the B allele frequent (BAF) and Log R ratio (LRT) is illustrated. **I-J**) Sequencing results for the validation of each patient specific LIS1 mutation with Sanger sequencing (P1, P2, P3, P5, P6, P7) or whole exome sequencing (P4). Scalebars: B-E: 50 µm; F-G: 20 µm.

**Supplementary Figure 2: LIS1-patient-cohort and control derived forebrain organoids. B)** Representative brightfield images of two control (C1, C2), two mild (P1, P2), 2 two moderate (P3, P4) and three severe (P5, P6, P7) LIS1 patient-derived organoids at day 20 and representative Hoechst staining of organoids derived from control or mild, moderate and severe grade patients at day 20±1. **B**) Quantification of structural VZ parameters in control and LIS1 patient-derived organoids at day 20. Central line in boxplot represents median, lower lines the 25^th^ and upper lines the 75^th^ percentile, Whiskers are 1.5 interquartile ranges. Individual dots represent the mean of one batch. Scale bar 200 µm

**Supplementary Figure 3: Characterization of cellular populations, developmental trajectories and proteomic networks in LIS1 patient-derived organoids. A)** Dot plot graph showing the expression of markers used to identify the different cellular populations (neuroepithelial cells (NE), cycling progenitors (CyP), radial glia cells (RG), intermediate progenitors (IP), transitory radial glia cells (t-RG), dorsal forebrain (dFB-N), ventral forebrain (vFB-N), midbrain (MB-N), interneurons (IN) and astroglial cells (G). **B**) UMAP plot showing the developmental trajectory of progenitor cells. **C)** Quantification of the percentage of neural progenitors (NES, CyP, RG, IP, t-RG) versus neurons and glial cells (dFB-N, vFB-N, MB-N, IN, G) in control, mild, moderate and severe conditions (left), and comparison between progenitor cell types in the four conditions (right), *p< 0.05, **p< 0.01, ***p< 0.001 **D)** Gene Ontology (GO) scores of apoptotic process and cell death for each sample. No difference in these processes is observed across severity grades. **E)** Heatmap depicting differentially regulated proteins between the mild, moderate, severe and control conditions (p<0.05, Wilcoxon’s r >0.75). Annotated are the unique top 10 differentially regulated proteins per condition. **F)** Progenitor cells derived from controls and LIS1 mutant lines stained for progenitor markers SOX2 and Nestin show a homogeneous induction of neuroectodermal fate (top). Withdrawal of factors from culture media result in premature differentiation, indicated by increased TUBB3 staining, consistent with the increased severity grade (center). Progenitor cells positive for caspase 3 do not increase across severity grades (bottom). **G)** String network of proteins included in the turquoise module. Proteins have been clustered by MCL clustering with the inflation parameter set to 3. Edges represent the degree of confidence for the interaction and only proteins forming complexes with a high degree of confidence (interaction score ≥ 0.7) have been plotted. Proteins in the red cluster are associated with protein catabolism, as they are either part of the proteasome or the COP9 signalosome. The yellow cluster includes proteins involved in cytoplasmic translation that are either structural components of the ribosome or enhance cytoplasmic translation. Green, blue, and purple clusters are associated with carbon metabolism, L-serine biosynthetic process and aminoacyl-tRNA ligase activity, respectively. **H)** emap plot of GO molecular functions overrepresented in differentially regulated genes and proteins, showing five categories, size of circles representing the number of differentially regulated features in GO term. Scale bars: F, top, bottom: 125 µm, middle: 100 µm.

**Supplementary Figure 4: Impact of epothilone D and CHIR99021 on cytoskeletal and morphological parameters in LIS1 patient-derived organoids. A)** Quantification of basal Ac-TUB strand density in DMSO and epothilone D (EpoD, 1nM) treated control and LIS1-patient derived organoids at day 15, statistical analyses in Sup. Table S13. **B)** Representative Ac-TUB images of control, mild, moderate and severe patient-derived organoids treated with DMSO or 1nM EpoD. Quantification of basal membrane length, apical membrane length, ventricle-like area, and total loop area, in DMSO or EpoD. **C)** Representative Hoechst staining of control, mild, moderate and severe LIS1-patient derived organoids treated with DMSO or EpoD. The yellow dotted lines indicate the edges of the VZ areas, the green dotted lines those of the cortical plate regions, and the green arrows illustrates the cortical plate thickness. **D)** Quantification of N-CAD diameter expansion in DMSO and EpoD treated control and LIS1-patient derived organoids at day 15, statistical analyses in Sup. Table S15. Central line in boxplot represents median, lower lines the 25^th^ and upper lines the 75^th^ percentile, Whiskers are 1.5 interquartile ranges. Individual dots represent each loop measurement. **E)** Representative N-cadherin (N-CAD) staining of mild, moderate and severe LIS1 patient-derived organoids treated with DMSO or EpoD. Yellow dotted lines define the VZ areas. **F)** VZ parameters of DMSO and EpoD or **G**) DMSO and CHIR treated control and patient-derived organoids at day 15. Quantification of apical and basal Ac-TUB strand density in DMSO or EpoD (1nM) treated control and LIS1 patient-derived organoids at day 15, statistics for (F) and (G) can be found in Supp. Table S14. Central line in boxplot represents median, lower lines the 25^th^ and upper lines the 75^th^ percentile, Whiskers are 1.5 interquartile ranges. Individual dots represent the mean of one batch (≥ 2 replicates) **H**) Representative DAPI images of control, mild, moderate and severe LIS1 patient-derived organoids treated with DMSO or CHIR99021 (CHIR, 1µM). The yellow dotted lines indicate the edges of the VZ areas, the blue dotted lines those of the cortical plate regions, and the blue arrows illustrates the cortical plate thickness. **I**) Quantification of cell division plane orientation in control and LIS1-patient derived organoids treated with DMSO or CHIR, statistical analyses in Sup. Table Error bars, ±SD. *P< 0.05, **P< 0.01, ***P< 0.001. Scalebars: B C, 200 µm; E, 20 µm, H, 200 µm.

