## Supplementary Figures for "Capturing the biology of disease severity in an organoid model of LIS1-lissencephaly"

A

| Patient | Age (years) | sex | Mutation | Dobyns grade |
|---|---|---|---|---|
| P1 mild | 8 | male | c.569-10 T>C splice site | 4-5 |
| P2 mild | 5 | male | c.569-10 T>C splice site | 4-5 |
| P3 moderate | 6 | female | c.13del frameshift | 3 |
| P4 moderate | 13 | female | delEx11 deletion | 3 |
| P5 severe | 4 | female | c.1002+1 G>A splice site | 1-2 |
| P6 severe | 18 | female | c.531 G>C missense | 2 |
| P7 severe | 3 | female | c.5445 C>T missense | 1-2 |

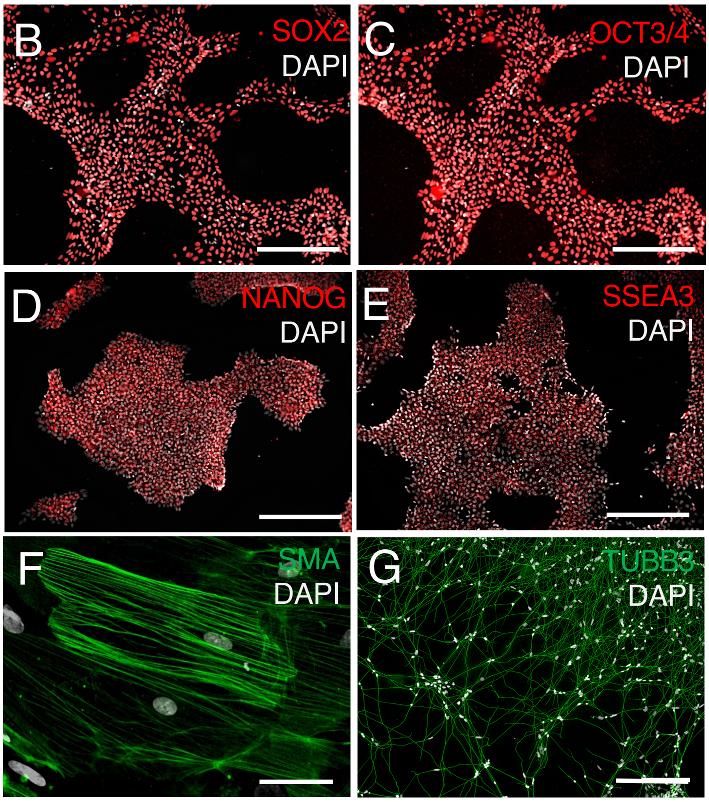


H

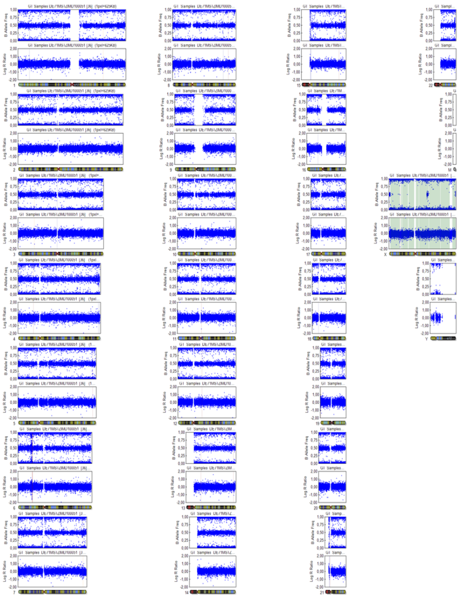

J

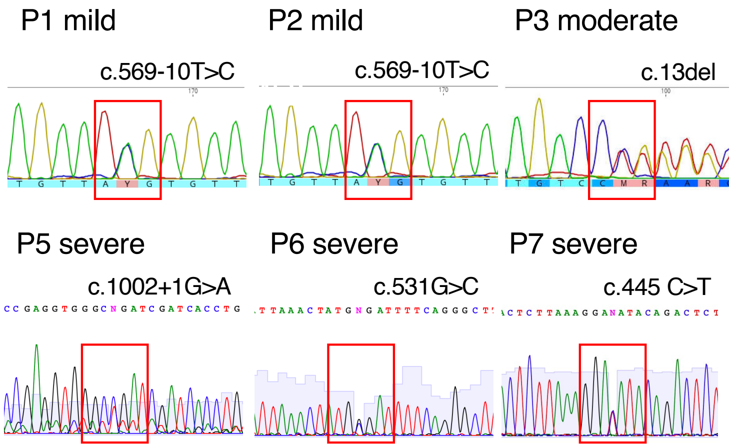


K

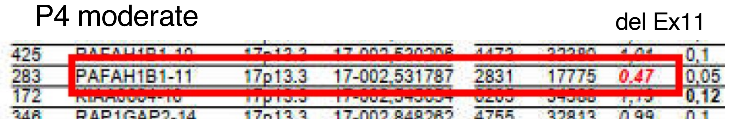

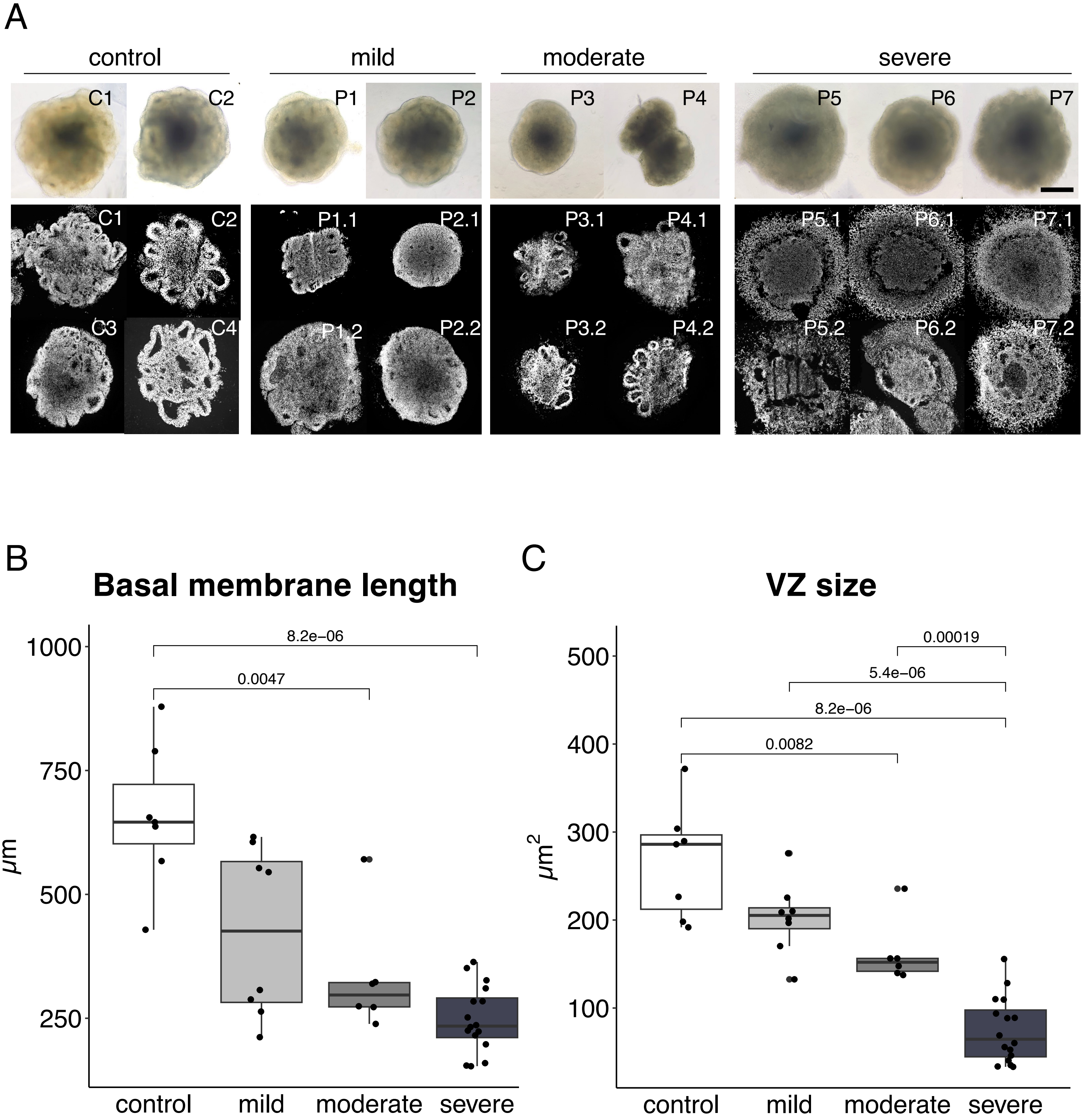
A
control
mild
moderate
severe
C1
C2
P1
P2
P3
P4
P5
P6
P7
C1
C2
C3
C4
P1.1
P2.1
P1.2
P2.2
P3.1
P4.1
P3.2
P4.2
P5.1
P6.1
P7.1
P5.2
P6.2
P7.2
B
Basal membrane length
μm
1000
750
500
250
8.2e-06
0.0047
control
mild
moderate
severe
C
VZ size
μm2
500
400
300
200
100
0.00019
5.4e-06
8.2e-06
0.0082
control
mild
moderate
severe

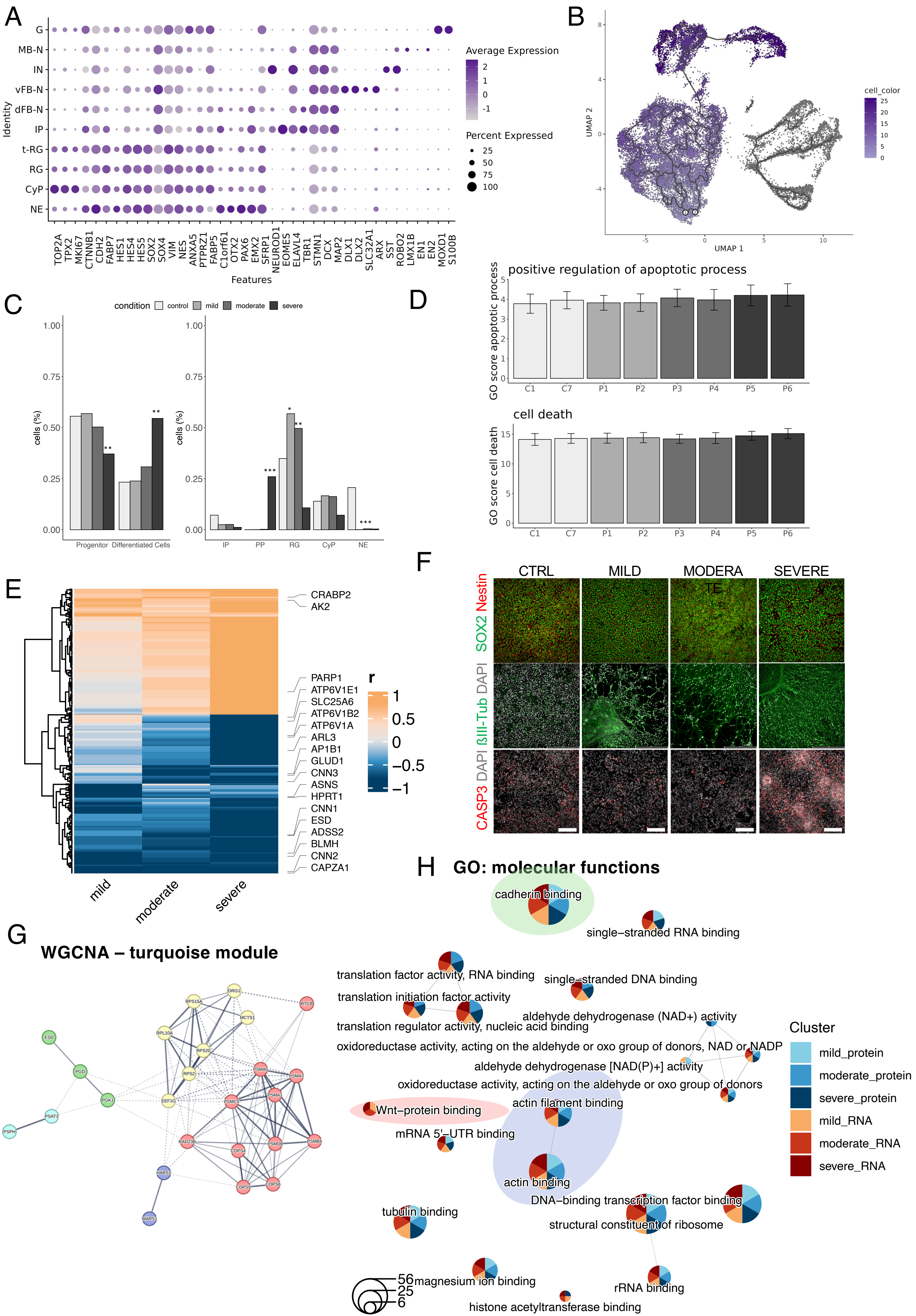

A
G
MB-N
IN
vFB-N
dFB-N
IP
t-RG
RG
CyP
NE
Identity
Average Expression
2
1
0
-1
Percent Expressed
25
50
75
100
TOP2A
TPX2
MKI67
CTNNB1
CDH2
FABP7
HES1
HES4
HES5
SOX2
SOX4
VIM
NES
ANXA5
PTPRZ1
FABP5
C1orf61
OTX2
PAX6
EMX2
SFRP1
NEUROD1
EOMES
ELAVL4
TBR1
STMN1
DCX
MAP2
DLX1
DLX2
SLC32A1
ARX
SST
ROBO2
LMX1B
EN1
EN2
MOXD1
S100B
Features
B
UMAP 2
UMAP 1
cell_color
25
20
15
10
5
0
C
condition
control
mild
moderate
severe
cells (%)
1.00
0.75
0.50
0.25
0.00
Progenitor
Differentiated Cells
IP
PP
RG
CyP
NE
D
positive regulation of apoptotic process
GO score apoptotic process
C1
C7
P1
P2
P3
P4
P5
P6
cell death
GO score cell death
E
CRABP2
AK2
PARP1
ATP6V1E1
SLC25A6
ATP6V1B2
ATP6V1A
ARL3
AP1B1
GLUD1
CNN3
ASNS
HPRT1
CNN1
ESD
ADSS2
BLMH
CNN2
CAPZA1
r
1
0.5
0
−0.5
−1
mild
moderate
severe
F
CTRL
MILD
MODERATE
SEVERE
SOX2 Nestin
ßIII-Tub DAPI
CASP3 DAPI
G
WGCNA – turquoise module
DRG1
RPS15A
RTCB
MCTS1
RPL10A
RPS20
ESD
PGD
RPS2
PSMA5
PSMA7
PGK1
EEF1G
PSMC5
PSMA1
PSAT1
PSPH
RAD23B
COPS4
PSMD8
PSMB4
HARS1
COPS5
COPS6
WARS1
H
GO: molecular functions
cadherin binding
single−stranded RNA binding
translation factor activity, RNA binding
single−stranded DNA binding
translation initiation factor activity
aldehyde dehydrogenase (NAD+) activity
translation regulator activity, nucleic acid binding
oxidoreductase activity, acting on the aldehyde or oxo group of donors, NAD or NADP
aldehyde dehydrogenase [NAD(P)+] activity
oxidoreductase activity, acting on the aldehyde or oxo group of donors
actin filament binding
Wnt−protein binding
mRNA 5'−UTR binding
actin binding
DNA−binding transcription factor binding
tubulin binding
structural constituent of ribosome
magnesium ion binding
rRNA binding
histone acetyltransferase binding
56
25
6
Cluster
mild_protein
moderate_protein
severe_protein
mild_RNA
moderate_RNA
severe_RNA

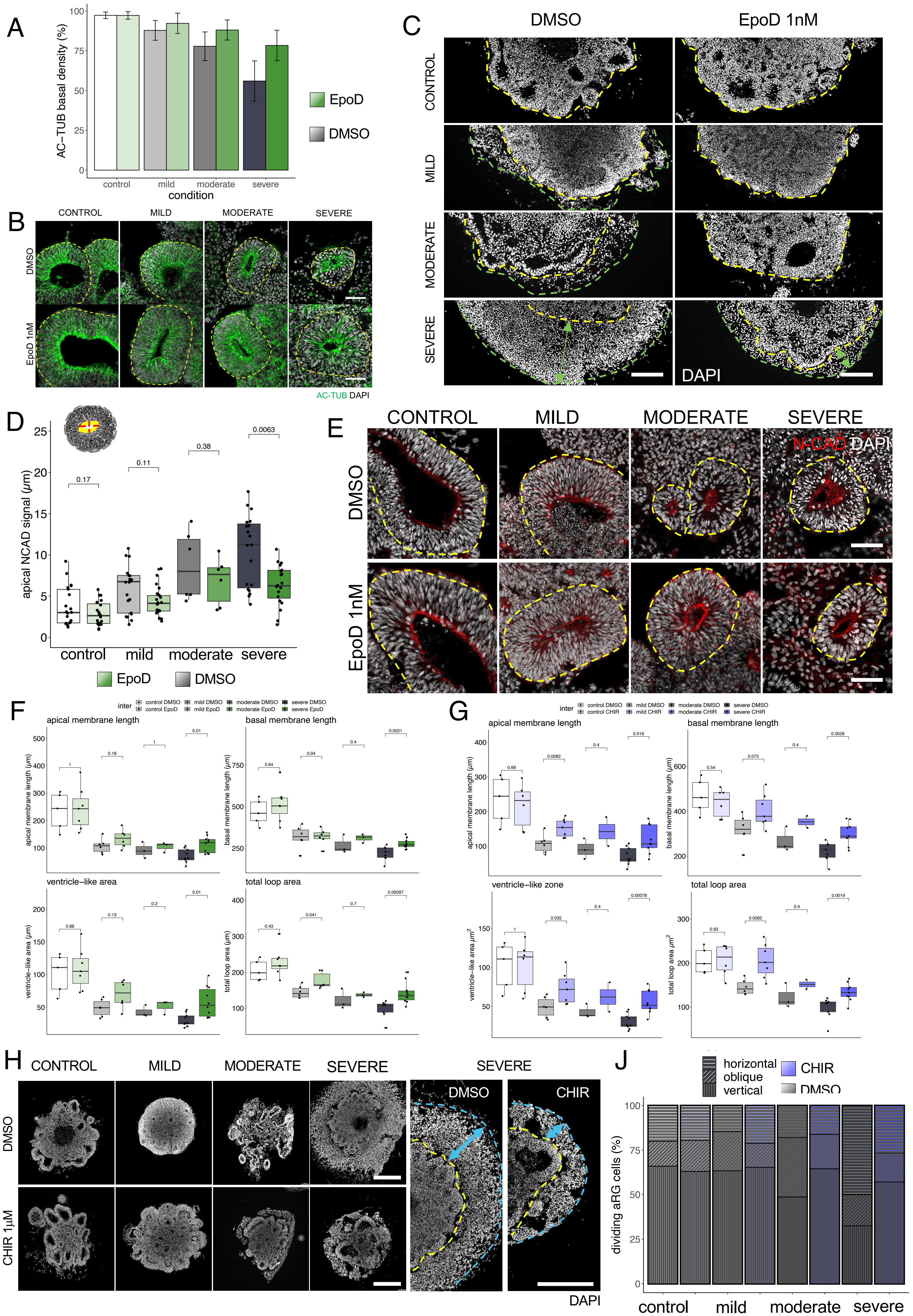

A
AC−TUB basal density (%)
control
mild
moderate
severe
condition
EpoD
DMSO
B
CONTROL
MILD
MODERATE
SEVERE
DMSO
EpoD 1nM
AC-TUB DAPI
C
DMSO
EpoD 1nM
CONTROL
MILD
MODERATE
SEVERE
DAPI
D
apical NCAD signal (μm)
0.17
0.11
0.38
0.0063
control
mild
moderate
severe
EpoD
DMSO
E
CONTROL
MILD
MODERATE
SEVERE
N-CAD DAPI
DMSO
EpoD 1nM
F
inter
control DMSO
control EpoD
mild DMSO
mild EpoD
moderate DMSO
moderate EpoD
severe DMSO
severe EpoD
apical membrane length
basal membrane length
ventricle−like area
total loop area
apical membrane length (μm)
basal membrane length (μm)
ventricle−like area (μm)
total loop area (μm)
G
control DMSO
control CHIR
mild DMSO
mild CHIR
moderate DMSO
moderate CHIR
severe DMSO
severe CHIR
ventricle−like zone
ventricle−like area μm²
total loop area μm²
H
CONTROL
MILD
MODERATE
SEVERE
DMSO
CHIR 1μM
CHIR
DAPI
J
horizontal
oblique
vertical
CHIR
DMSO
dividing aRG cells (%)
control
mild
moderate
severe
